# Bcl-xL interaction with VDAC1 reduces mitochondrial Ca^2+^ uptake, allowing the establishment of Therapy-Induced Senescence

**DOI:** 10.64898/2025.12.05.692660

**Authors:** Andrea Puebla-Huerta, Pablo Morgado-Cáceres, Camila Quezada- Gutierez, César Casanova-Canelo, Roberto Rosales-Rojas, José Alberto López-Domínguez, Inés Marin, Sandra A. Niño, Pablo Cruz, Micheel Saldivia-Piña, Sergio Linsambarth, Osman Díaz- Rivera, Oscar Cerda, Ariela Vergara-Jaque, Manuel Serrano, Christian Gonzalez-Billault, Ulises Ahumada-Castro, J. César Cárdenas

**Affiliations:** Center for Integrative Biology, Faculty of Sciences, Universidad Mayor, Santiago 8580745, Chile; Geroscience Center for Brain Health and Metabolism, Santiago 8580745, Chile; Center for Bioinformatics, Simulation and Modeling, Faculty of Engineering, Universidad de Talca, Talca, Chile, 3460000; Centro de Investigación del Cáncer, CSIC-University of Salamanca, Campus Unamuno, 37007, Salamanca, Spain; Department of Biochemistry and Molecular Biology, University of Salamanca, Campus Unamuno, 37007, Salamanca, Spain; Department of Immunology, Genentech, South San Francisco, USA; Department of Biology, Laboratory of Cellular and Neuronal Dynamics, Faculty of Sciences, Universidad de Chile, Santiago, Región Metropolitana 7800003, Chile; Millennium Nucleus of Ion Channel-Associated Diseases (MiNICAD), Santiago 8380453, Chile; Program of Cellular and Molecular Biology, Institute of Biomedical Sciences (ICBM), Faculty of Medicine, Universidad de Chile, Santiago 8380453, Chile; Cambridge Institute of Science, Altos Labs, Granta Park, Cambridge CB21 6GP, UK; Department of Neuroscience, Faculty of Medicine, Universidad de Chile, Chile; Public Health Unit, Institute for Nutrition and Food Technology (INTA), Universidad de Chile, Chile; The Buck Institute for Research on Aging, Novato, USA; Department of Chemistry and Biochemistry, University of California Santa Barbara, Santa Barbara, CA, 93106, USA; Department of Neuroscience and Physiology, Upstate Medical University, Syracuse, NY, 132010, USA

**Author notes:** Senior author and to whom correspondence should be addressed: Cesar Cardenas, PhD Center for Integrative Biology Camino la Piramide 5750 Universidad Mayor Santiago, CHILE, Ulises Ahumada-Castro, PhD Facultad de Odontología Bellavista 7, Recoleta Universidad San Sebastián Santiago, Chile. Present address: Facultad de Odontología y Ciencias de la Rehabilitación, Universidad San Sebastián, Bellavista, Santiago, Chile (Ulises Ahumada-Castro).

## Abstract

Cellular senescence, a state of irreversible growth arrest, is characterized by various phenotypic changes, including altered mitochondrial function. While the role of mitochondria in senescence is well-established, the mechanisms underlying their involvement remain unclear. Here, we investigate the early stages of therapy-induced senescence (TIS) and identify a novel anti-apoptotic mechanism mediated by Bcl-xL and VDAC1, two key regulators of mitochondrial calcium (Ca²⁺) homeostasis.

We find that Bcl-xL expression increases in early TIS cells and localizes to the mitochondria, where it interacts with the voltage-dependent anion channel 1 (VDAC1). This interaction dampens mitochondrial Ca²⁺ uptake, thereby preventing Ca²⁺ overload and apoptosis. Disrupting this interaction using the BH3 mimetic ABT-263 or Bcl-xL-targeting siRNA increases mitochondrial Ca²⁺ uptake, leading to apoptosis and blocking the formation of senescent cells.

These findings uncover a previously unrecognized role of the Bcl-xL–VDAC1 axis in regulating mitochondrial Ca²⁺ dynamics during the onset of senescence. Our work provides mechanistic insight into how senescent cells evade apoptosis,highlighting potential therapeutic targets for selectively eliminating them in cancer and age-related diseases.

## INTRODUCTION

Cellular senescence, in its canonical form, is defined by a permanent and irreversible arrest of cell proliferation, accompanied by distinctive phenotypic changes such as altered secretory profiles (senescence-associated secretory phenotype, SASP), activation of tumor suppressor pathways, chromatin remodeling, genomic instability, and organelle dysfunction^1^. Several stimuli can trigger senescence. For instance, excessive cellular replication leads to replicative senescence, primarily driven by telomere shortening and activation of the DNA damage response (DDR)^2^. Activation of oncogenes—including Ras, BRAF, AKT, E2F1, and cyclin E—or the loss of tumor suppressors such as PTEN and NF1, can initiate oncogene-induced senescence (OIS), a critical tumor-suppressive mechanism^3^. Additionally, therapy-induced senescence (TIS) can arise as a result of sublethal damage caused by chemotherapeutic agents. TIS is characterized by a delayed onset, typically manifesting several days after drug exposure, and shares many features with other forms of senescence, including growth arrest and SASP^4^.

The senescent phenotype is defined by a combination of distinct cellular and molecular features. A hallmark of senescent cells is the persistent activation of the DDR, which is often marked by increased phosphorylation of histone H2A.X (γ-H2A.X) and elevated levels of p53 protein^5^. Chromatin remodeling in conjunction with chronic DDR also promotes the upregulation of Cyclin-Dependent Kinase inhibitors (CDKis), particularly p16INK4a (hereafter p16) and p21CIP1 (hereafter p21), encoded by the CDKN2A and CDKN1A loci, respectively³. Another defining characteristic of senescent cells is their elevated expression of anti-apoptotic proteins, which confers certain resistance to apoptosis and contributes to their persistence within tissues^6,7^. This resistance is mediated by members of the B-cell lymphoma 2 (Bcl-2) protein family—key regulators that integrate survival and apoptotic signals depending on their expression, subcellular localization, and binding partners^8,9^. Notably, the anti-apoptotic protein Bcl-xL is frequently upregulated in senescent cells, with regulation occurring at both transcriptional and translational levels. Importantly, pharmacological inhibition of Bcl-xL has been shown to selectively induce cell death in senescent cells, a phenomenon known as senolysis^10–14^.

In addition to the upregulation of CDK inhibitors and resistance to apoptosis, cellular senescence is marked by profound structural and functional changes in organelles. Among these, mitochondrial alterations are particularly prominent. A substantial body of evidence has demonstrated that mitochondrial dysfunction is both a hallmark and a driver of senescence^15–17^. There are two noteworthy points in this regard. First, mitochondria have been identified as potential targets of senolytic drugs, signifying their crucial role in the survival of senescent cells. Second, and most intriguingly, the senescent phenotype cannot be achieved without the involvement of mitochondria, emphasizing their indispensable role in its development^15–17^. Senescent cells exhibit several mitochondrial anomalies, including changes in membrane potential, impaired electron transport chain activity, altered oxidative phosphorylation efficiency, and disruptions in cellular bioenergetics^18^. Of particular interest is the dysregulation of mitochondrial calcium (Ca²⁺) homeostasis^19,20^. Senescent cells display increased mitochondrial Ca²⁺ uptake, driven by enhanced Ca²⁺ transfer from the endoplasmic reticulum (ER), alongside changes in mitochondrial membrane potential, NAD⁺/NADH and AMP/ATP ratios, and mitochondrial morphology^21^. Despite this Ca²⁺ overload, senescent cells often maintain viability. However, excessive and unregulated mitochondrial Ca²⁺ can trigger apoptosis via outer mitochondrial membrane (OMM) permeabilization and the release of cytochrome c^22–24^.

The mechanisms enabling senescent cells to maintain their balance and survive with excessive mitochondrial Ca^2+^ remain poorly understood. In cancer cells, the anti-apoptotic protein Bcl-xL has been shown to interact with the voltage-dependent anion channel 1 (VDAC1) on the OMM, limiting mitochondrial Ca²⁺ influx and preventing apoptosis^8,25,26^. Whether a similar mechanism operates in therapy-induced senescence (TIS) has not yet been explored.

In this study, we identify an early and critical molecular event required for the establishment of TIS. Specifically, we show that the interaction between Bcl-xL and VDAC1, which occurs early after chemotherapeutic exposure, limits Ca²⁺ transfer from the ER to mitochondria, thereby preventing apoptotic cell death and allowing senescence to proceed. These findings uncover a key molecular determinant in the balance between senescence and apoptosis,highlighting a potential target for novel senotherapeutic strategies.

## RESULTS

To systematically examine whether anti-apoptotic signaling pathways are activated during the early stages of senescence induction, we treated proliferating human diploid IMR90 fibroblasts with 250 nM doxorubicin (Doxo) for 24 hours, a concentration previously shown to induce a stable senescent phenotype by day 10^27^. Remarkably, as early as 24 hours post-treatment, we detected increased levels of p21, p53, and γ-H2A.X, all canonical markers of DNA damage response and early senescence (Figure 1A). These findings confirm that molecular features associated with the senescent program are rapidly activated in response to chemotherapeutic challenge.

**Figure 1:**
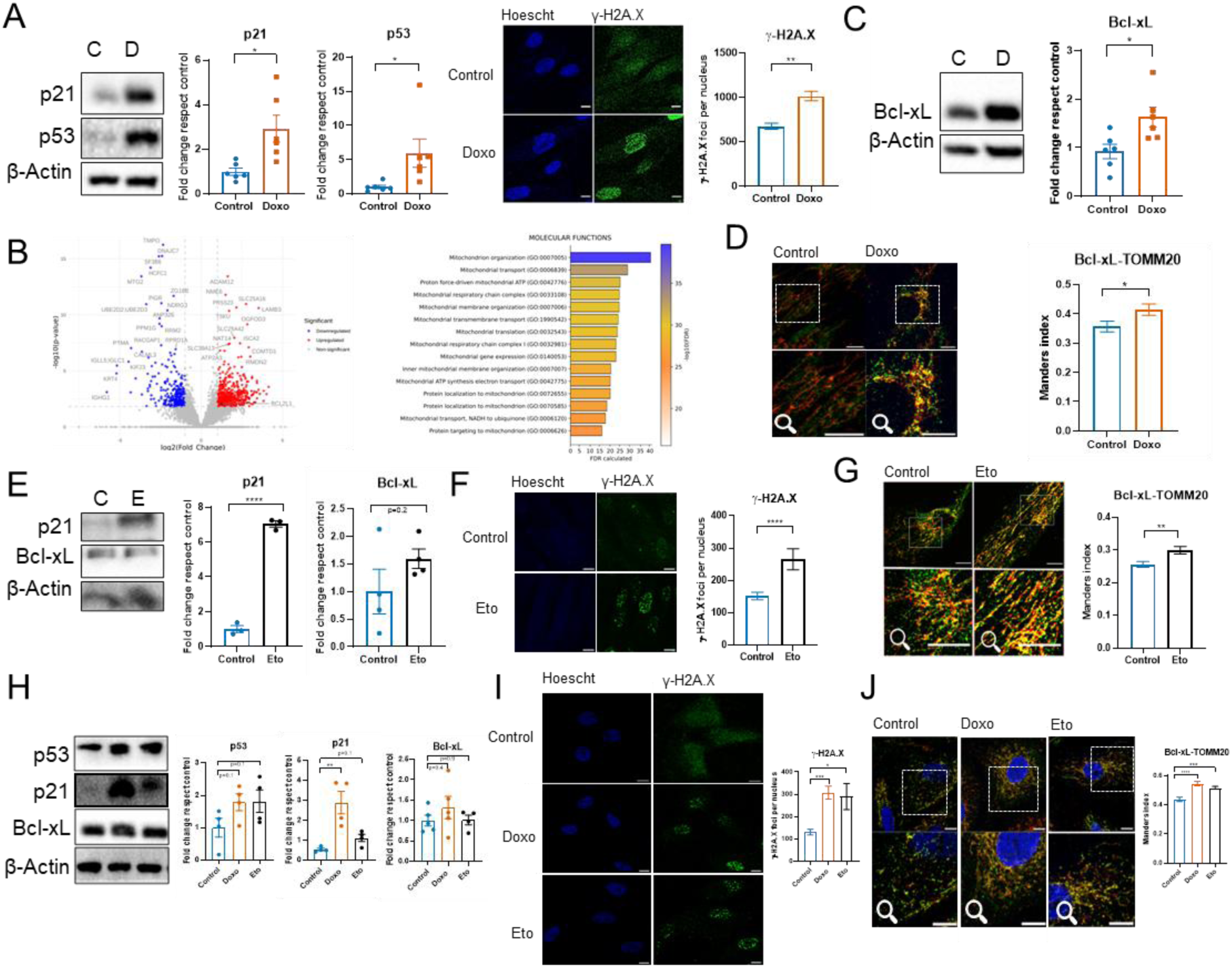
Early senescence is associated with increased Bcl-xL expression and mitochondrial localization. (A) Induction of early senescence in IMR90 fibroblasts by treatment with doxorubicin for 24 hours. Western blot analysis showing increased levels of p21, p53, and immunofluorescence of γ-H2A.X (938 ± 37 foci) respect (619 ± 28 foci); markers of early senescence. (B) Differential expression and functional enrichment analysis. The volcano plot (left) shows significantly upregulated (red) and downregulated (blue) genes in early senescent cells; notably, Bcl2L1 (Bcl-xL) is among the most strongly upregulated genes. The bar plot (right) displays enriched molecular functions of the upregulated genes, primarily associated with mitochondrial organization and regulation. (C) Western blot analysis confirming increased Bcl-xL protein levels in early senescent cells. (D) Immunofluorescence staining showing increased colocalization of Bcl-xL (green) with the mitochondrial marker TOMM20 (red) in early senescent cells (0,43 ± 0,02) respect (0,35 ± 0,02). (E), (F), (G) Similar to doxorubicin, treatment with etoposide (40 nM) also induced increased senescence markers p21, p53, γ-H2A.X (388 ± 23 foci) respect (159 ± 13 foci); Bcl-xL expression and mitochondrial localization in IMR90 cells (0,3 ± 0,012) respect (0,256 ± 0,009). (H), (I), (J) Increased senescence markers p21, p53, γ-H2A.X (308 ± 29 foci) in doxorubicin and etoposide (154 ± 31 foci) respect (131 ± 13); Bcl-xL upregulation and mitochondrial localization were also observed in A549 cell lines treated with doxorubicin (0,546 ± 0,016) and etoposide (0,516 ± 0,011) respect (0,439 ± 0,016). All graphs shown average ±SEM. N=6 for western blot, C: control, D: doxorubicin, E: etoposide. N=3 for immunofluorescence, ∼50 cells/N. * p<0,05, ** p<0,01, *** p<0,001 and **** p<0,0001. Manders’ coefficient was used to quantify colocalization. Scale bars=10 µm. Doxorubicin (250 nM) and etoposide (40 nM). Panels 1A–1G: T-Test; panels 1H–1J: one-way ANOVA.

Given that anti-apoptotic proteins are known to localize to and anchor within mitochondrial membranes^28–30^, we performed a comprehensive proteomic analysis of cellular membrane fractions under the conditions described above, to assess whether Bcl-xL shows increased membrane association. Our analysis revealed that Bcl2L1 (Bcl-xL) was among the most strongly upregulated genes in early senescent cells (Figure 1B, left). Gene Ontology (GO) enrichment analysis further indicated that upregulated genes were primarily associated with mitochondrial organization and regulation (Figure 1B, right). These analyses revealed that Bcl-xL was the predominant anti-apoptotic protein enriched in the mitochondrial membrane compartment. Together, these results suggest a selective recruitment or stabilization of Bcl- xL at mitochondrial membranes during the early stages of TIS.

Whole-cell protein analysis by Western blot confirmed an increase in Bcl-xL expression, showing a 1.6-fold elevation compared to control conditions (Figure 1C).^10,13,14^. Bcl-xL can be found in the mitochondria, ER, and the cytosol^31^. Interestingly, during early stages TIS, we observed an increased mitochondrial localization of Bcl-xL, as evidenced by its enhanced colocalization with the mitochondrial marker TOMM20 (Manders’ coefficient: Control = 0.35 ± 0.02; Doxo = 0.41 ± 0.02) (Figure 1D, S1A).

To assess whether the observed increase in Bcl-xL expression and its mitochondrial localization represents a general response to chemotherapeutic stress, we extended our analysis to an additional senescence-inducing agent. We evaluated the effects of etoposide (Eto), a topoisomerase II inhibitor widely used in TIS models^32^. Similar to Doxo, a 24-hour treatment with Eto (40 μM) resulted in a significant upregulation of senescence-associated markers, including p21 (Figure 1E) and γ-H2A.X (Figure 1F). Although Bcl-xl protein levels did not change significantly after Eto exposure (Figure 1E), its localization in the mitochondria increased, as evidenced by enhanced colocalization with the mitochondrial outer membrane marker TOMM20 (Manders’ coefficient: Control = 0.26 ± 0.01; Eto = 0.30 ± 0.01) (Figure 1G, S1B). Next, we investigated whether the observed increase in Bcl- xL expression and mitochondrial localization also occurs in other cell types. To this end, we examined the human lung carcinoma cell line A549, which is widely used in studies of cellular senescence^25^. Consistent with our findings in IMR90 fibroblasts, treatment with either Doxo or Eto tended to induce classical senescence markers in A549 cells, including increased levels of p21, p53 (Figure 1H), and γ-H2A.X (Figure 1I). Although the increase in Bcl-xL protein levels was less pronounced in A549 cells compared to IMR90 fibroblasts, its mitochondrial localization was notably enhanced, as evidenced by increased colocalization with the mitochondrial marker TOMM20 (Manders’ coefficient: Control = 0.439 ± 0.02; Doxo = 0.546 ± 0.02; Eto = 0.516 ± 0.01) (Figure 1J and S1C).

Bcl-xL has been reported to exert a range of anti-apoptotic effects through diverse mechanisms of action^33^. One such mechanism, which has not yet been explored in the context of the senescent phenotype, involves the interaction between Bcl-xL and the voltage-dependent anion channel 1 (VDAC1) at the mitochondrial outer membrane. This interaction has been shown to reduce mitochondrial Ca²⁺ uptake, thereby preventing Ca²⁺ overload and apoptosis^8,25,33^. Given our observation of increased Bcl-xL accumulation in mitochondria during early TIS, we sought to determine whether Bcl-xL interacts with VDAC1. To this end, we employed a Proximity Ligation Assay (PLA). As shown in Figure 2A, early TIS cells exhibited a significant increase in PLA signal (Bcl-xL–VDAC1 interaction dots: Control = 154 ± 10.8, Doxo = 278 ± 18.8 dots/cell), indicating enhanced interaction between Bcl-xL and VDAC1. Consistent with these results, *in silico* docking analysis revealed that Bcl-xL binds to VDAC1 through coordinated interactions between its BH domains and the VDAC1 β-barrel. In particular, the BH1 and BH2 domains constitute the main contact sites, forming electrostatic interactions with polar residues at the binding interface between the two proteins (Figure 2B).

**Figure 2:**
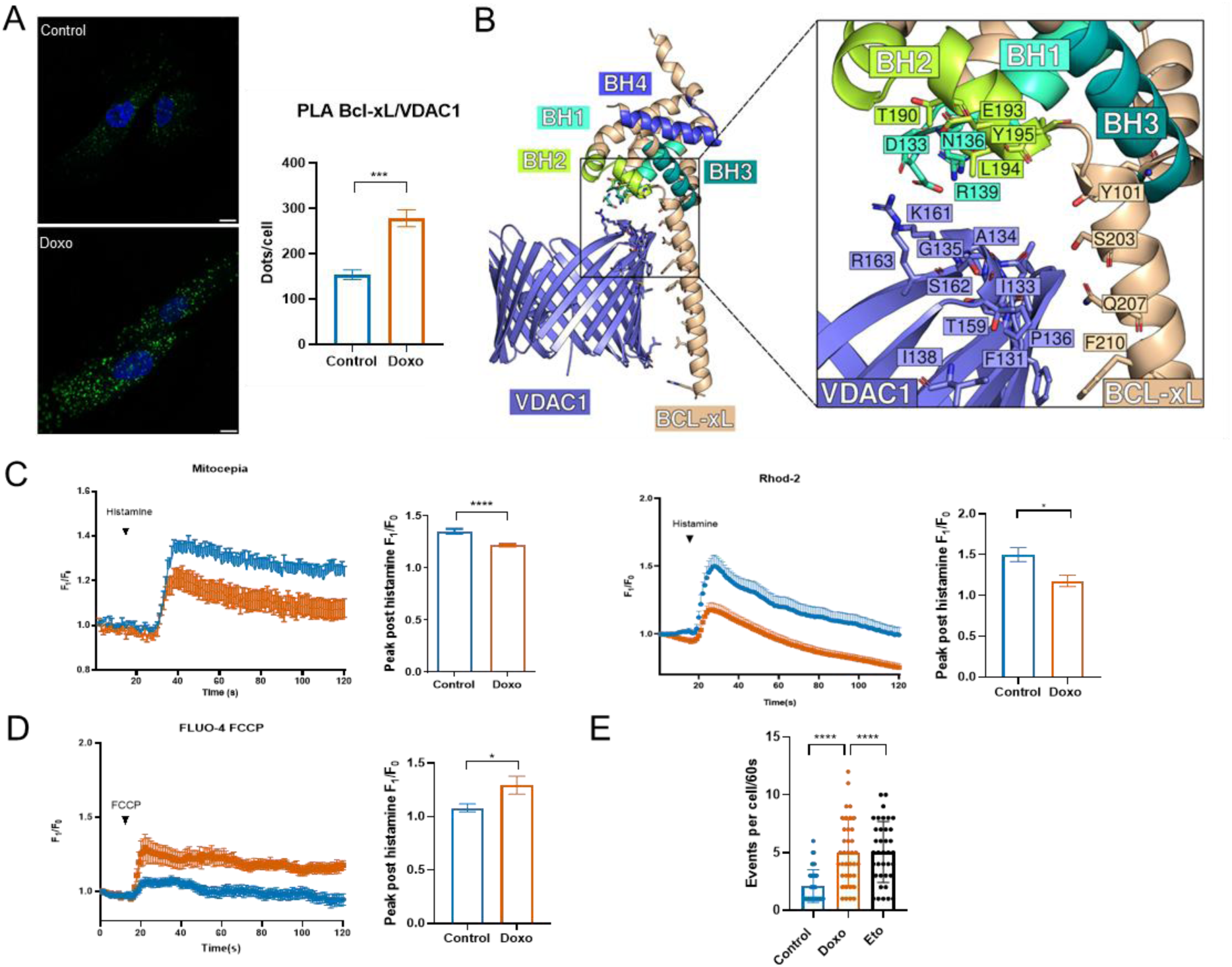
Bcl-xL interacts with VDAC1 to regulate mitochondrial Ca^2+^ uptake during early senescence. (A) Proximity ligation assay (PLA) showing increased interaction between Bcl-xL and VDAC1 in early senescent cells. Green dots indicate protein-protein interactions in doxorubicin-treated cells (298 ± 23) compared with control cells (205 ± 18). Graphs shown average ± SEM. N=3. ∼10 cells/N and *** p<0,001. Scale bar=10 µm. (B) Structural docking model of the Bcl-xL–VDAC1 interaction. The BH1-BH4 domains of Bcl-xL are highlighted in distinct colors. The close-up view depicts the interaction interface, highlighting key residues from both Bcl-xL and VDAC1 that contribute to stabilizing the complex. (C) Mitochondrial Ca^2+^ uptake is decreased in early senescent cells compared to control cells, measured using Rhod-2 AM in doxorubicin (1,178 ± 0,069) respect (1,500 ± 0,085) and CEPIA2mt probes in doxorubicin (1,219 ± 0,062) respect (1,352 ± 0,102). N=3. ∼20 cells/N. Histamine 300 µM. (D) Levels of mitochondrial Ca^2+^ are increased in early senescent cells compared to control cells, measured using Fluo-4 in doxorubicin (1,294 ± 0,084) respect (1,081 ± 0,036). ∼20 cells/N. FCCP 1 µM. * p<0,05 and **** p<0,0001. (E) Spontaneous Ca^2+^ release events recorded in IMR90 cells one hour after being stimulated with either 250nM doxorubicin (Doxo), 40 nM etoposide (Eto) or vehicle (control) in normal media (N=3). ****p < 0.001. See also Video S1, S2 and S3. All panels: T-Test.

As previously mentioned, the interaction between Bcl-xL and VDAC1 has been shown to reduce mitochondrial Ca²⁺ uptake^25^. To determine whether this mechanism is active during TIS, we evaluated mitochondrial Ca²⁺ uptake in IMR90 cells following a 24- hour treatment with Doxo. Mitochondrial Ca²⁺ levels were measured using both Rhod-2 AM and CEPIA2mt probes after stimulation with 300 μM histamine to induce IP3R-mediated Ca²⁺ release. As expected, early TIS cells exhibited a significant reduction in mitochondrial Ca²⁺ uptake compared to control cells (Figure 2C). Importantly, IP3R-mediated Ca²⁺ release, assessed by Fluo-4 AM, remained intact (Figure S2A), as did ER luminal Ca²⁺ levels measured using G-CEPIAer (Figure S2B). Interestingly, a significant reduction in ER Ca²⁺ refilling was observed in senescent cells relative to controls, indicating extensive rewiring of intracellular Ca^2+^ homeostasis associated with the senescent state (Figure S2C). The diminished mitochondrial Ca^2+^ uptake was not associated with changes in the protein levels of VDAC1, MCU, MICU1, type 1, 2 and 3 IP3R (Figure S2D).

A similar reduction in mitochondrial Ca²⁺ uptake was also observed in early TIS IMR90 cells induced with Eto (Figure S2E), as well as a comparable trend in A549 cells treated with either Doxo or Eto (Figure S2F).

In established late senescent cells, an accumulation of Ca²⁺ within the mitochondrial matrix has been previously reported^34–37^. To determine whether this phenomenon occurs early during the onset of senescence, we assessed mitochondrial Ca²⁺ content in IMR90 cells treated with Doxo for 24 hours. Cells were challenged with the mitochondrial uncoupler FCCP in the presence of the cytosolic Ca²⁺ indicator Fluo-4 AM to monitor mitochondrial Ca²⁺ release. As shown in Figure 2D, early TIS cells exhibited greater mitochondrial Ca²⁺ accumulation compared to controls. These findings suggest that the observed reduction in mitochondrial Ca²⁺ uptake in early TIS may represent a compensatory response to mitochondrial Ca²⁺ overload.

Notably, Doxo and eto treatment did not trigger acute cytosolic Ca²⁺ mobilization, as assessed by Fluo-4 AM (Figure S2G). However, when we examined spontaneous IP3R- mediated Ca²⁺ release events one hour after Doxo or Eto exposure, these events were significantly more frequent than in untreated cells (Figure 2E and S2H). We propose that these enhanced and frequent IP3R-driven Ca²⁺ transients may contribute to the mitochondrial Ca²⁺ overload observed early (24 hours after Doxo) and the establishment of cellular senescence.

Altogether, our findings indicate that during the first 24 hours of TIS onset, there is a significant increase in spontaneous IP3R-mediated Ca²⁺ release events, accompanied by elevated mitochondrial matrix Ca²⁺ content. Concurrently, Bcl-xL expression is upregulated and predominantly localized to the mitochondria, where it interacts with VDAC1 via its BH1 and BH2 domains. This interaction likely contributes to the observed reduction in mitochondrial Ca²⁺ uptake, serving as a regulatory mechanism to prevent mitochondrial Ca²⁺ overload during early senescence.

### BH-3 mimetics can avoid VDAC-Bcl-xL interaction in the early steps of TIS

To assess whether the interaction between Bcl-xL and VDAC1 contributes to the reduced mitochondrial Ca²⁺ uptake observed in early TIS cells, we aimed to disrupt this interaction pharmacologically. Using a structural comparison between the molecular docking of the VDAC1–Bcl-xL complex and the previously resolved Bcl-xL–ABT-263 complex (PDB ID: 4QNQ), we identified a homologous binding site of high interest for targeted inhibition. ABT-263 is a specific BH3 mimetic inhibitor that binds with high affinity to both Bcl-xL and Bcl-w^14,38^. Our structural analysis showed that ABT-263 partially occupies a binding pocket in Bcl-xL that overlaps with the interface used for VDAC1 binding. As illustrated in Figure 3A, ABT-263 (yellow) engages several residues within the BH1-BH3 domains of Bcl-xL. When superimposed with the VDAC1–Bcl-xL docking model, ABT-263 binding coincides with key residues in the BH1 and BH2 domains (e.g., D133, N136, R139, L194, and Y195) that are also critical for VDAC1 association. These findings suggest that ABT-263 may compete with VDAC1 for binding to Bcl-xL, thereby potentially disrupting the Bcl-xL–VDAC1 interaction.

**Figure 3:**
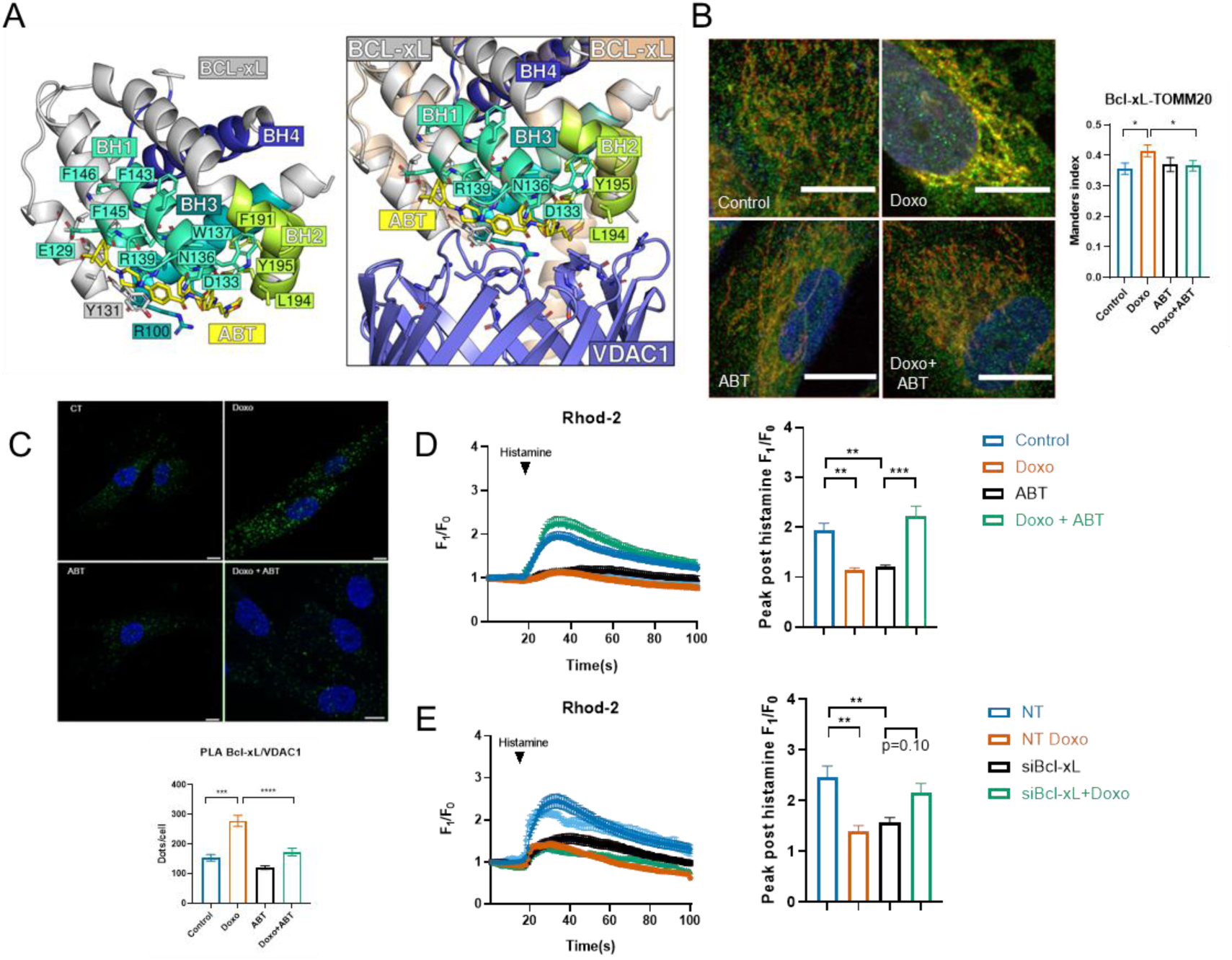
Bcl-xL interacts with VDAC1 to regulate mitochondrial Ca^2+^ uptake during early senescence. (A) Crystal structure of the Bcl-xL–ABT-263 complex showing ABT-263 (yellow) bound to the BH1-BH3 domains. Superimposition with the VDAC1–Bcl-xL model reveals overlapping binding sites, suggesting that ABT-263 may competitively interfere with VDAC1 association. (B) Immunofluorescence staining showing decreased colocalization of Bcl-xL (green) with TOMM20 (red) in early senescent cells treated with ABT-263. (C) Proximity ligation assay (PLA) demonstrating a reduction in the interaction between Bcl-xL and VDAC1 in early senescent cells treated with ABT-263. (D) Mitochondrial Ca^2+^ uptake, measured using Rhod-2 AM and CEPIA2mt probes, is restored in early senescent cells treated with ABT-263. (E) siRNA-mediated knockdown of Bcl-xL, but not other Bcl-2 family members, reverses the impaired mitochondrial Ca^2+^ uptake in early senescent cells. All panels: one-way ANOVA. All panels: one-way ANOVA; all other panels: T-Test.

Based on our structural analysis, we hypothesized that ABT-263 treatment would disrupt the Bcl-xL–VDAC1 interaction observed during early TIS (Figure 2A), therefore reducing Bcl-xL mitochondrial localization. Consistent with this hypothesis, immunofluorescence analysis revealed a marked decrease in the co-localization of Bcl-xL with the mitochondrial marker TOMM20 in the presence of ABT-263 (Manders’ coefficient: Control = 0.35 ± 0.02; Doxo = 0.41 ± 0.02; ABT = 0.37± 0.02; Doxo+ABT= 0.36 ± 0.01) (Figures 3B and S3A). Furthermore, PLA demonstrated a significant reduction in the interaction between Bcl-xL and VDAC1 following treatment with ABT-263(Figure 3C). Interestingly, ABT-263 also led to a decrease in total Bcl-xL protein levels (Figure S3B).

Given this loss of mitochondrial localization and interaction, we next evaluated whether ABT-263 affected mitochondrial Ca²⁺ uptake in early TIS cells. Notably, ABT-263 restored mitochondrial Ca^2+^ uptake in early TIS cells to levels comparable to those observed in untreated control cells (Figure 3D), supporting the notion that mitochondrial Bcl-xL inhibits Ca²⁺ uptake via interaction with VDAC1. Interestingly, in non-senescent (non-TIS) cells, ABT-263 reduced mitochondrial Ca^2+^ uptake and slowed the kinetics of the Ca^2+^ signal (Figure 3D), indicating a context-dependent effect of Bcl-xL on mitochondrial Ca^2+^ dynamics. Notably, ABT-263 did not alter IP3R-mediated cytosolic Ca²⁺ release (Figure S3B), nor did it affect the expression levels of key components of the mitochondrial calcium uptake machinery, including MICU1 and MCU, or the type 1, 2, and 3 IP3R isoforms (Figure S3C). To confirm the inhibition of Bcl-xL, mediates the effect of ABT-263, we performed a Bcl-xL siRNA knockdown. As shown in Figure 3E, Bcl-xL levels were reduced by approximately 90% after 24 hours of siRNA incubation, and this knockdown persisted for at least 48 hours (Figure S4C). Based on this, we added doxorubicin after the initial 24-hour siRNA incubation, to mimic the combined effect of ABT-263. Figure 3E shows that only the combination of Bcl-xL siRNA and doxorubicin was able to reverse the diminished mitochondrial Ca²⁺ uptake, recapitulating the effect observed with ABT-263 treatment.

These findings demonstrate that during the early stages of senescence development, Bcl-xL interacts with VDAC1 at the mitochondrial membrane, leading to a reduction in mitochondrial Ca²⁺ uptake. This effect can be reversed either by pharmacological inhibition of Bcl-xL using ABT-263, a mechanism not previously described, or by genetic knockdown of Bcl-xL, highlighting a novel regulatory role for Bcl-xL in mitochondrial Ca^2+^ homeostasis during the onset of the senescent phenotype.

### VDAC-Bcl-xL interaction is essential to the viability of TIS cells in its early steps

Our results indicate that Bcl-xL interacts with VDAC1 at the onset of therapy-induced senescence (TIS), contributing to the observed reduction in mitochondrial Ca²⁺ uptake, a phenomenon previously associated with resistance to apoptosis^22,39–42^. To investigate whether this interaction confers an anti-apoptotic advantage during early TIS, we utilized ABT-263, which disrupts the Bcl-xL–VDAC1 interaction and restores mitochondrial Ca²⁺ uptake. Cell death was assessed using SYTOX staining. As shown in Figure 4A, ABT-263 treatment led to a nearly threefold increase in the percentage of SYTOX-positive (dying) cells compared to both untreated controls and Doxo-treated cells alone. Similarly, knockdown of Bcl-xL also induced significant cell death (Figure 4B).

**Figure 4:**
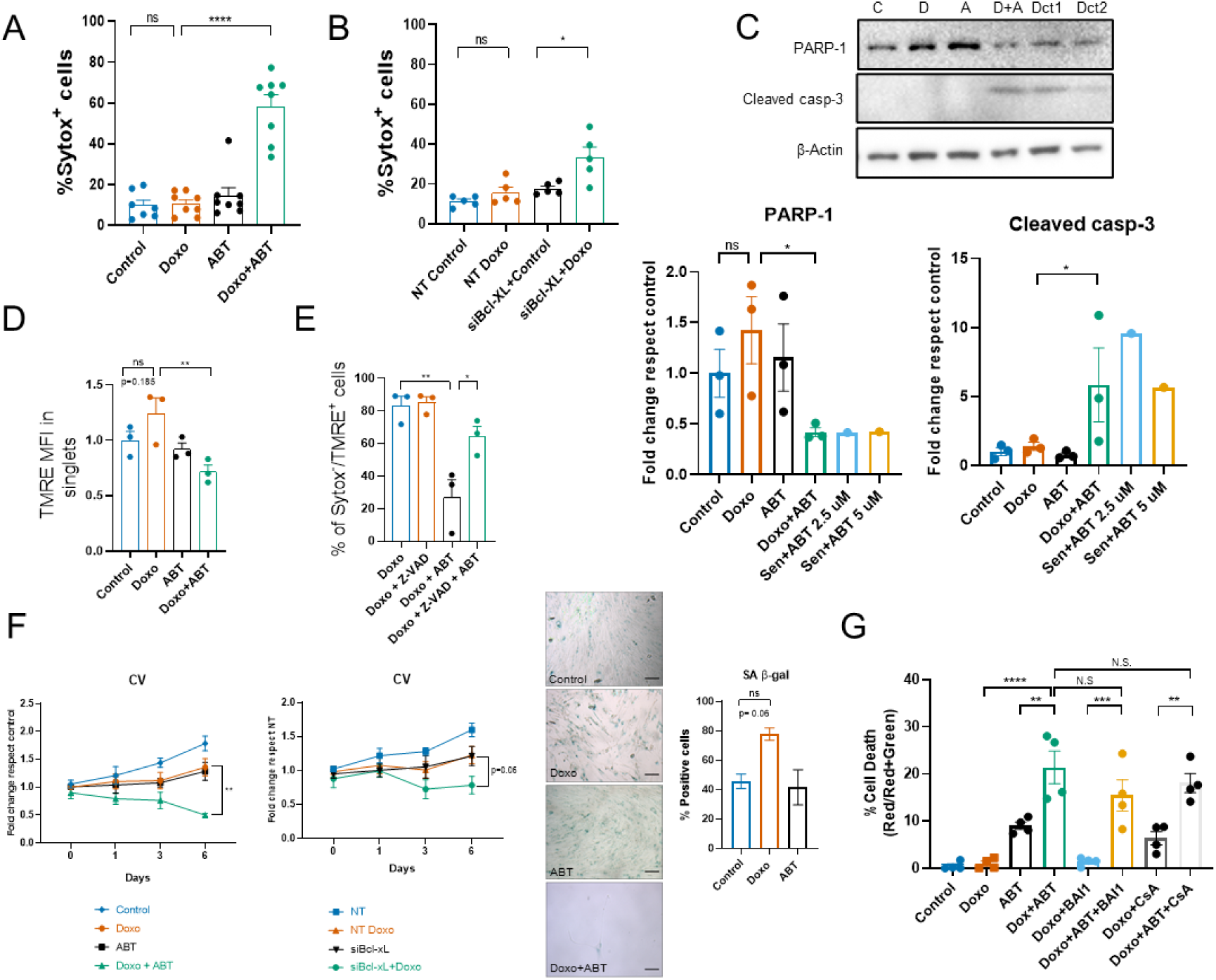
**Disruption of Bcl-xL–VDAC1 interaction triggers apoptosis in early senescent cells**. (A) ABT-263 (ABT, 3 µM) increases cell death in early TIS (24 h post-doxorubicin, Doxo 250 nM), quantified as SYTOX Green+ cells. (B) siRNA-mediated Bcl-xL knockdown similarly increases SYTOX Green+ cells in early TIS. (C) ABT-263 promotes apoptosis, evidenced by increased cleaved PARP-1 and cleaved caspase-3 by immunoblot; Dct1 and Dct2 are death-control conditions in 12-day senescent cells treated with ABT-263 (2.5 and 5 µM). (D) Mitochondrial membrane potential (Δψm), assessed by TMRE, decreases upon ABT-263 in the presence of Doxo, consistent with mitochondrial depolarization during apoptosis. (E) The pan-caspase inhibitor Z-VAD-FMK rescues ABT-263+Doxo-induced cell death, indicating caspase-dependent apoptosis. (F) Time-course viability/proliferation (crystal violet) shows modest effects of ABT-263 alone, growth arrest with Doxo, and a marked loss of viability with Doxo+ABT; analogous reductions in viability are observed with siBcl-xL ± Doxo. SA-β-galactosidase staining at endpoint confirms that Doxo+ABT prevents the establishment of the senescent phenotype, mirroring the viability curves. (G) Inhibition of Bcl-xL (ABT-263) increases mitochondrial Ca2+ uptake and cell death that are not rescued by the mPTP inhibitor cyclosporin A or the BAX/BAK pore inhibitor BAI1. G: growing; D: doxorubicin (250 nM); A: ABT-263 (3 µM); D+A: Doxo+ABT-263. Dct1: 12-day senescent cells + ABT-263 (2.5 µM); Dct2: 12-day senescent cells + ABT-263 (5 µM). Data:mean ± SEM. (n=3 independent experiments). Statistics: one-way ANOVA; *P<0.05.

This pro-apoptotic effect of ABT-263 was not limited to Doxo-induced TIS. A comparable increase in cell death was observed when Eto was used to induce TIS in IMR90 cells treated with ABT-263 (Figure S4A), as well as in A549 cells exposed to either Doxo or Eto in combination with ABT-263 (Figure S4B). It is important to note that ABT-263 alone also increased cell death in non-senescent cells, as previously reported^32,43^, consistent with its known pro-apoptotic activity.

To determine whether the cell death observed upon ABT-263 treatment at the onset of TIS corresponds to apoptosis, we assessed PARP-1 levels as an indirect indicator of PARP-1 cleavage, along with the presence of cleaved caspase-3. Notably, ABT-263 treatment at the initiation of TIS resulted in a marked reduction in full-length PARP-1 and a concomitant increase in cleaved caspase-3, consistent with the activation of apoptotic pathways (Figure 4C). Additionally, a significant loss of mitochondrial membrane potential— an early hallmark of apoptosis—was observed in ABT-263-treated cells (Figure 4D). This effect was prevented by the pan-caspase inhibitor Z-VAD-FMK, further confirming the induction of apoptosis (Figure 4E). Significantly, the presence of ABT-263 at the initiation of TIS completely abrogated the establishment of the senescent phenotype at later time points (Figure 4F), highlighting a critical window in which Bcl-xL-mediated mitochondrial protection supports senescence survival and suggests a potential therapeutic strategy for senescence clearance.

Mitochondrial Ca²⁺ overload is a well-known trigger of apoptosis, typically mediated through the opening of the mitochondrial permeability transition pore (mPTP) or the formation of BAX/BAK-dependent pores^44–46^. To determine whether either of these pathways contributes to the cell death observed in early TIS cells treated with ABT-263, we employed pharmacological inhibitors: Cyclosporin A (CsA) to block mPTP opening, and BAI1 to inhibit BAX/BAK pore formation. As shown in Figure 4G, neither CsA nor BAI1 prevented cell death under these conditions, suggesting that apoptosis is mediated by an alternative, mPTP- and BAX/BAK-independent mechanism.

### BH3-mimetics in mice prevents the accumulation of senescent cells after treatment with doxorubicin

To evaluate the physiological relevance of our findings, we assessed whether ABT- 263 could prevent the development of the senescent phenotype *in vivo*. Mice were divided into three experimental groups: (1) Doxo, receiving a single intraperitoneal injection of doxorubicin (10 mg/kg); (2) Doxo + ABT-263, receiving the same single dose of Doxo followed by five consecutive daily injections of ABT-263; and (3) Control, receiving 10% DMSO (Doxo vehicle) and corn oil (ABT-263 vehicle) for five consecutive days (Figure 5A). Mice were euthanized 8 days after Doxo administration (and 4 days after the final ABT-263 injection), and tissues were harvested for analysis. The lung has been shown to be particularly sensitive to senescence-inducing stimuli^47–49^. RT-qPCR analysis of lung tissue revealed increased expression of p21 variant 1 (p21v1), p21 variant 2 (p21v2), and IL-6 mRNAs in the Doxo-treated group, consistent with the induction of a senescent phenotype. Strikingly, these transcript levels were restored to near-control values in the Doxo + ABT- 263 group (Figure 5B), supporting the senescence-suppressive effect of ABT-263 *in vivo*^38^. In agreement, γ-H2A.X immunofluorescence staining of lung sections showed an increased number of DNA damage–positive cells in Doxo-treated mice, which was significantly reduced upon ABT-263 treatment (Figure 5C), further confirming inhibition of TIS induction. Finally, to determine whether ABT-263 promotes apoptosis *in vivo*, we assessed cleaved caspase-3 and cleaved PARP as markers of apoptosis. Doxo treatment alone modestly increased the number of cleaved caspase-3–positive cells and cleaved PARP levels in lung tissue (Figure 5D and E). However, the combination of Doxo and ABT-263 resulted in a substantially greater increase in both markers, indicating enhanced apoptotic cell death. Together, these data suggest that ABT-263 suppresses the development of TIS *in vivo* by promoting apoptosis.

**Figure 5.**
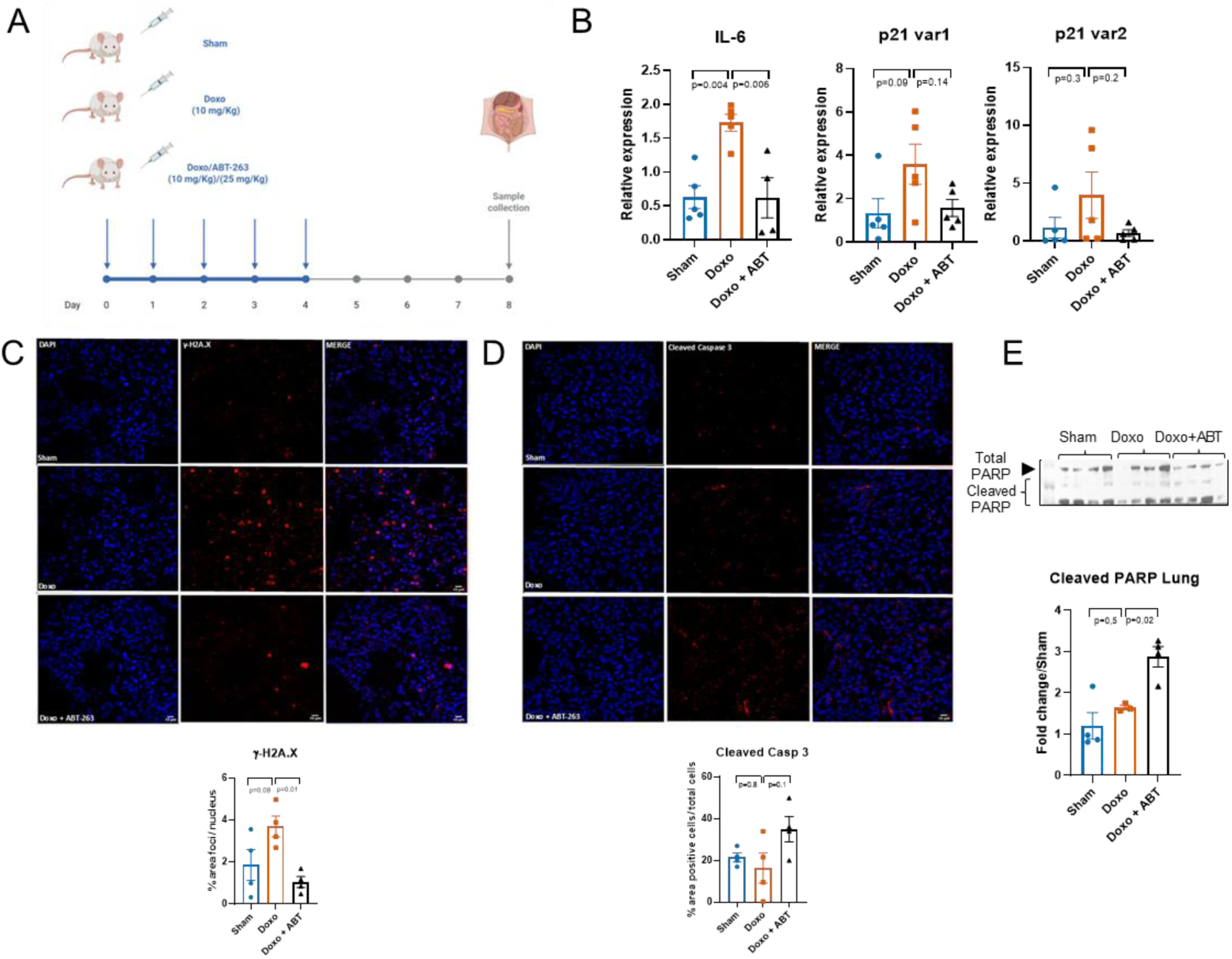
ABT-263 promotes apoptosis and prevents senescence in vivo. (A) Experimental design: Mice were treated with a single intraperitoneal dose of doxorubicin (Doxo, 10 mg/kg), ABT-263 (25 mg/kg), a combination of both (Doxo + ABT), or vehicle only (Sham). Lungs were collected on day 7 for downstream analysis. (B) RT-qPCR analysis of lung tissue revealed increased expression of senescence-associated genes following Doxo treatment, including IL-6 (Sham: 0.6300 ± 0.3774; Doxo: 1.732 ± 0.2823; Doxo + ABT: 0.6200 ± 0.5957), p21 var 1 (Sham: 1.336 ± 1.514; Doxo: 3.596 ± 2.070; Doxo + ABT: 1.574 ± 0.8890), and p21 var 2 (Sham: 1.156 ± 1.978; Doxo: 3.968 ± 4.501; Doxo + ABT: 0.6920 ± 0.6920). Co-treatment with ABT-263 reversed the Doxo-induced increase in all three transcripts. Five animals were used per group; one outlier was excluded from the IL-6 dataset based on statistical criteria. (C) Immunofluorescence staining for γ-H2A.X (red) and nuclear counterstaining with DAPI (blue) showed increased DNA damage (as % of nuclear area positive for γ-H2A.X) in Doxo-treated lungs (Sham: 1.856 ± 0.3774; Doxo: 3.689 ± 0.4937), which was significantly reduced with ABT-263 co-treatment (Doxo + ABT: 1.046 ± 0.2561). Four animals were analyzed per group. (D) Cleaved caspase-3 (red) staining revealed reduced apoptosis in Doxo-treated lungs compared to Sham (Sham: 21.50 ± 2.102; Doxo: 16.40 ± 7.297), while co-treatment with ABT-263 led to a significant increase in cleaved caspase-3-positive cells (Doxo + ABT: 35.00 ± 6.137). Values represent percentage of cleaved caspase-3-positive cells over total nuclei (n=4 per group). (E) Western blot analysis of cleaved PARP levels in lung homogenates showed enhanced apoptosis in the Doxo + ABT group (Sham: 1.203 ± 0.3219; Doxo: 1.647 ± 0.05389; Doxo + ABT: 2.876 ± 0.2457; fold change relative to Sham, normalized to β-actin). Four animals were used per group. one-way ANOVA followed by appropriate post hoc tests for all comparisons. Figure 5A created with BioRender.com. All panels: one-way ANOVA.

In summary, our findings demonstrate that during the first 24 hours of senescence development, referred to here as early TIS, there is an increase in Bcl-xL expression and its localization to the mitochondria. At the mitochondria, Bcl-xL interacts with VDAC1, suppressing mitochondrial Ca²⁺ uptake and thereby preventing mitochondrial Ca²⁺ overload and apoptosis. Disruption of this interaction, either by siRNA-mediated knockdown of Bcl-xL or pharmacological inhibition using the BH3 mimetic ABT-263, restores mitochondrial Ca²⁺ uptake, promotes Ca²⁺ overload, and induces apoptosis. This apoptotic response effectively prevents the establishment of senescent cells both *in vitro* and *in vivo* (Figure 6).

**Figure 6.**
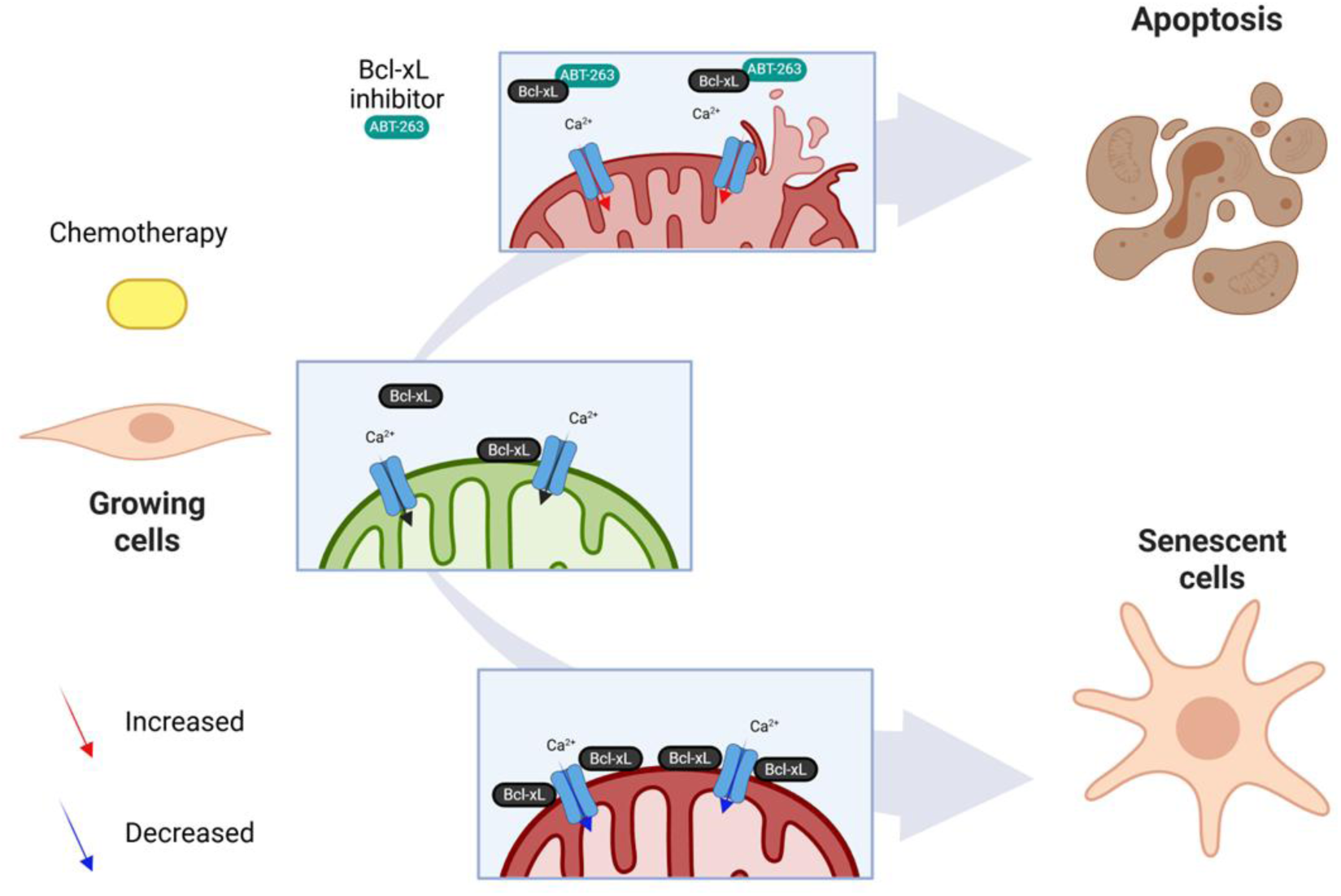
**Proposed model: BCL-xL regulates mitochondrial Ca²**⁺ **homeostasis to determine cell fate upon chemotherapy.** Under normal proliferative conditions, BCL-xL interacts with the outer mitochondrial membrane channel VDAC1 to limit mitochondrial Ca²⁺ uptake, thereby preventing Ca²⁺ overload and apoptosis. Following chemotherapy, this regulatory interaction is maintained, favoring the induction of senescence through reduced mitochondrial Ca²⁺ influx. However, pharmacological inhibition of BCL-xL with ABT-263 disrupts this interaction, leading to increased mitochondrial Ca²⁺ accumulation and triggering apoptosis in damaged or senescent cells. Red arrows indicate increased Ca²⁺ uptake; blue arrows indicate reduced uptake. Schematic created with BioRender.com.

## DISCUSSION

We described that Bcl-xL protein levels increase in the early hours of TIS cells, localizing in the mitochondria and interacting with VDAC1, which decreases the mitochondrial Ca^2+^ uptake.All these effects can be avoided using ABT-263, a BH3 mimetic,which decreases Bcl-xL protein levels and interrupts the Bcl-xL–VDAC1interaction. Interestingly, ABT-263 changes Ca^2+^ fluxes inside the TIS cells during the early hours, particularly avoiding the decreased mitochondrial Ca^2+^ uptake. As we mentioned, ABT-263 has been described as a senolytic drug^10,13,14^ but the mechanisms of how it induces cell death, particularly apoptosis, remain unknown. Considering this information along with our data, we propose that cells treated with chemotherapeutic drugs face a dichotomous fate. On the one hand, if Bcl-xL protein levels increase, localizing in the mitochondria and interacting with VDAC1, they prevent a mitochondrial Ca^2+^ overload in the early hours of TIS cells. On the other hand, the interruption of the Bcl-xL–VDAC1 interaction, using ABT-263 or siRNA against Bcl-xL, cells enter cell death, particularly apoptosis (Figure 3A).

The increase in Bcl-xL and its localization to the mitochondria in early TIS cells, as demonstrated by our proximity ligation assays and (Figure 2A and 2C), aligns with previous observations of the role of BCL-xL in cancer cell survival. However, our study extends this understanding by showing that the Bcl-xL–VDAC1 interaction specifically contributes to the regulation of mitochondrial Ca^2+^ uptake in early TIS. This is significant because mitochondrial Ca^2+^ overload is a known trigger for apoptosis through mechanisms such as the opening of the mitochondrial permeability transition pore (mPTP). By restricting Ca^2+^ entry into the mitochondria, the interaction between Bcl-xL and VDAC1 mitigates this apoptotic signal, promoting the survival of cells undergoing senescence.

Our data further reveal that disrupting the Bcl-xL–VDAC1 interaction using the BH3 mimetic ABT-263 or siRNA targeting Bcl-xL reverses this protective mechanism. This disruption leads to increased mitochondrial Ca²⁺ uptake and apoptosis, highlighting the anti-apoptotic role of the Bcl-xL–VDAC1 interaction in early TIS. Importantly, this apoptosis was not prevented by Cyclosporin A (CsA) or Bax Channel Blocker, indicating that it occurs independently of mPTP opening and BAX/BAK pore formation. These findings suggest that alternative mitochondrial mechanisms drive apoptosis upon disruption of Bcl-xL–VDAC1. Supporting this notion, a study by Heimer et al. demonstrated that the small molecule Raptinal can induce rapid apoptosis through mitochondrial outer membrane permeabilization (MOMP) and cytochrome c release, independent of BAX, BAK, or BOK^50^. Similarly, Chen et al. recently reported that certain Bcl-2 family inhibitors can induce apoptosis through VDAC1 oligomerization, bypassing BAX/BAK dependency^51^. This suggests the existence of alternative pathways that facilitate MOMP and apoptosis without the involvement of these pro-apoptotic proteins. Therefore, the ability of ABT-263 to disrupt the Bcl-xL–VDAC1 interaction and induce apoptosis points to its potential as a senolytic agent capable of targeting early senescent cells, thereby preventing their accumulation The observation of a selective reduction in mitochondrial Ca^2+^ uptake adds another layer of complexity to the Ca^2+^ signaling alterations that occur during senescence. It suggests that early senescence is associated with a unique modulation of Ca^2+^ homeostasis, which may be critical for the shift towards a senescent phenotype. This regulation of Ca^2+^ dynamics by Bcl-xL may be a key determinant in the balance between cell survival and death during the early stages of senescence induction.

Our *in vivo* studies using a mouse model of senescence further emphasize the relevance of our findings. Treatment with ABT-263 in combination with Doxo reduced the expression of senescence-associated markers such as p21 and IL-6 in lung tissues, as well as the accumulation of γ-H2A.X-positive cells. This suggests that targeting the Bcl-xL–VDAC1 interaction could be a viable strategy to limit the persistence of senescent cells in tissues exposed to chemotherapeutic stress. Moreover, the induction of apoptosis in ABT-263- treated mice, as evidenced by increased cleaved caspase-3 and PARP, supports the notion that disrupting this interaction triggers apoptotic pathways, thereby counteracting the survival mechanisms in TIS cells.

Our results provide critical insights into the mechanisms underlying the survival of TIS cells, revealing a novel interaction between Bcl-xL and VDAC1 that modulates mitochondrial Ca^2+^ dynamics. This interaction represents a potential therapeutic target for the selective elimination of senescent cells, particularly during the early stages of senescence when they are most vulnerable to interventions that disrupt their pro-survival adaptations. Future studies could explore the broader implications of targeting Bcl-xL–VDAC1 interactions in other models of senescence and assess the long-term effects of BH3 mimetics such as ABT- 263 on tissue homeostasis and aging.

In conclusion, the identification of the Bcl-xL–VDAC1 interaction as a regulator of mitochondrial Ca^2+^ homeostasis in early TIS provides a new understanding of the cellular adaptations that enable the survival of senescent cells. These findings open avenues for the development of targeted therapies that could mitigate the deleterious effects of senescent cell accumulation in aging and age-related diseases.

## ACKNOWLEDGMENTS

We thank the Unidad de Microscopía Avanzada, Pontificia Universidad Católica de Chile and the Unidad de Microscopía Universidad Mayor, UM2i for their support during imaging.

## MATERIALS AND METHODS

### Reagents

From *Thermo Fisher Scientific* (Waltham, Massachusetts, USA): Dulbeccós modified Eagle medium-high glucose (DMEM-HG) (12100-046), DMEM-HG no glutamine and calcium (21068028), trypsin (0.25%)-EDTA phenol red (25200072), antibiotic-antimycotic (100X) (15240062), trypan Blue solution 0.4% (15250061), Opti-MEM™ GlutaMAX™ supplemented (51985034), DMEM FluoroBrite™ (A1896701), Fluo-4/AM (F-14201), Rhod2/AM (R-1245MP), Lipofectamine™ 3000 (L3000015), Lipofectamine™ RNAiMAX (13778150), MitoTracker™ Green FM (M7514), TMRE (T669), Hoechst 33342 (H3570), Fluoromount-G mounting medium (00-4958-02), Pierce™ Bradford plus protein assay (23236), SuperSignal™ West Pico PLUS chemiluminescent substrate (34580), PageRuler™ Plus prestained protein ladder (26619), Image-iT™ fixative solution (FB002), X-Gal (B-1690), SYTOX™ blue (s11348). From *Cytiva* (Chicago, Illinois, USA): Fetal bovine serum (HC.SV30160.03), Amersham Hybond P 0.45 PVDF blotting membrane (10600023). From *Sigma-Aldrich* (St. Louis, Missouri, USA): Histamine dihydrochloride (H7250-5G), PLA Duolink® (DUO92014-100RXN), crystal violet (C6158), oligomycin (O4876), carbonyl cyanide 4-(trifluoromethoxy) phenylhydrazone (FCCP) (C2920), antimycin A (A8674), rotenone (R8875), protease Inhibitor Cocktail (P2714), PhosSTOP™ (4906837001). From *Tocris Bioscience* (Bristol, United Kingdom): From *Merck* (Darmstadt, Alemania): Acrylamide bis-acrylamide 29:1, 40% solution (1690-OP), triton® X-100 (112298), methanol (106009), 10X PBS (6505-OP), dimethyl sulfoxide (317275), hydrochloric acid fuming 37% (100317), sodium hydroxide (106498), TWEEN® 20 (655205), RIPA buffer (20-188). From *Corning Life Sciences* (New York, USA): From *Agilent Technologies* (Santa Clara, CA, USA): Seahorse XF calibrant solution (100840-000), seahorse XFe96 FluxPak (102601-100). From *Santa Cruz Biotechnology* (Dallas, Texas, EE. UU.): BSA (sc-2323), Z-VAD-FMK (Z-VAD) (sc-3067). From *Winkler* (Santiago, RM, Chile): TRIS buffer (BM-0585), glycerol (BM- 0800), 2-mercaptoethanol (BM-1200), SDS (BM-1750), bromophenol blue (AZ-0395), glycine (BM-0820), methanol (AL-0210), glutaraldehyde (WK- 106), NaCl (SO-1455), KCl (PO-1260), Tween-20 (016520).

*<u>Plasmids.</u>* From *Addgene* (Watertown, MA, USA.): pCMV CEPIA2mt (58218).

*<u>siRNAs.</u>* From *Thermo Fisher Scientific* (Waltham, Massachusetts, USA): siRNA BCL2L1 (4390824), siRNA BCL2L2 (4390824), Non target (AM4641).

*<u>Antibodies.</u>*From *Cell Signaling Technology* (Danvers, MA, USA): anti-IP_3_R1 (8568), anti MCU (14997), anti-p21 Waf1/Cip1 (12D1) (2947), anti-PARP1 (46D11), anti-P53 (9282S), anti-Bcl-xL(54H6), anti-MCU (D2Z3B), anti-MICU1 (D4P8Q). From *Abcam* (Cambridge, UK): anti-VDAC1 (AB10527), anti-VDAC1 (AB14734), anti-TOMM20 (AB56783), anti-cleaved caspasa-3 (AB32042). From *Becton & Dickinson* (Franklin Lakes, NJ, USA.): anti IP_3_R3 (610313). From *Thermo Fisher Scientific* (Waltham, Massachusetts, USA): Anti-IP_3_R2 (PA1-904), anti β-tubulin (32-2600), anti-mouse secondary antibody Alexa Fluor™ 350 (A11045), anti-rabbit secondary antibody Alexa Fluor™ 488 (A11008), anti-mouse secondary antibody Alexa Fluor™ 488 (A11001), anti-rabbit secondary antibody Alexa Fluor™ 568 (A11011), anti-mouse secondary antibody Alexa Fluor™ 568 (A11004), anti-rabbit secondary antibody Alexa Fluor™ 647 (A21244), anti-mouse secondary antibody Alexa Fluor™ 647 (A21235), anti-rabbit secondary antibody HRP (31460), anti-mouse secondary antibody HRP (31430). From *Santa Cruz Biotechnology* (Dallas, Texas, EE. UU.): anti-β-actin (sc-47778), anti-HSP90 (13119sc). From *Millipore* (Burlington, Massachusetts, EE. UU): Anti-phospho-Histone H2A.X (Ser139) (05-636).

### Cell Culture, treatment, and detection of SA-β-Gal

IMR90 (ATCC® CCL186™) cell line was maintained in Dulbecco’s modified Eagle’s medium, supplemented with 10% fetal bovine serum (FBS; *Biological Industries*), 100 U/ml penicillin, 100 mg/ml streptomycin, and 0.25 µg/ml amphotericin B (antibiotic-antimycotic solution), at 37 °C, 5% CO_2_ and 3% O_2_. To induce early senescence, cells were incubated with 250 nm doxorubicin for 24 h, the day after seeding the cells. In late senescence experiments (6 days/12 days), cells were incubated with 250 nm doxorubicin for 48 h and the culture medium was refreshed every 2 days. SA-β-Gal activity was evaluated as previously described^52^. All cell counts were performed manually by using ImageJ and performed in at least three independent replicates. From each replicate, 100 cells were counted. All experiments, both control and doxorubicin treated cells, were conducted using between 5,000 and 10,000 cells per cm^2^.

### Plasmid and siRNAs transfection

For *siRNA*-based transient silencing experiments, transfections were performed using Lipofectamine RNAiMax (Thermo Fisher Scientific) as transfecting reagent, according to the manufacturer’s instructions. Briefly, 1.5x10^5^ cells/well were seeded in 6-well plates and grown for 24 h. For transfections, a mix was prepared by diluting siRNAs (60 pmol/well) and Lipofectamine RNAiMax (15 µL /well) in Opti-MEM, preparing a total volume of 300 µL/well. This mixture was incubated for 15 minutes to allow the formation of complexes, and then it was dropped into each well containing the cells in complete medium. After 12 h the culture media were replaced with fresh medium and cells were incubated at 37°C in a 5% CO_2_ and 3% O_2_ incubator. The experiments were performed 24-48 h post-transfection as indicated in the figures. In the same way, *plasmid transfection* was performed using Lipofectamine™ 3000 (Thermo Fisher Scientific) on cells plated in 6-well plate. The transfection mixture was prepared considering 2 µg/well of plasmid DNA, 5 µL P3000, and 10 µL Lipofectamine™ 3000 diluted in Opti-MEM. After 12 h the culture media were replaced with fresh medium and cells were incubated at 37°C in a 5% CO_2_ and 3% O_2_ incubator. The experiments were performed 24-48 h post-transfection.

### Cell viability and mitochondrial membrane potential

Viability assays and determination of mitochondrial membrane potential were assessed using flow cytometry (FACSCantoA™; Becton & Dickinson™) by SYTOX blue (Max. ex/em: 444/480 nm) staining and TMRE (Max. ex/em: 549/574 nm), respectively. For these experiments, cells were seeded in 6-well plates, grown for 24 h, and then treated with different stimuli. To determine probe incorporation as a reflection of loss of cell viability in each experimental condition, culture medium was collected and cells trypsinized, pelleted, and lastly stained with 30 nM SYTOX blue or 5 nM TMRE at room temperature for 30 minutes in PBS, protected from light. For the analysis of cell death by apoptosis, cells were pre-incubated with the pan-caspase inhibitor Z-VAD at 40 µM. Each sample was analyzed by cytometry, counting in each one a total of 30,000 events. The resulting data was analyzed using the software “Cyflogic” version *1.2.1*.

### Cell proliferation assay

Cell proliferation after treatments was assessed at 0, 1, 3, and 6 days by staining with crystal violet. Cells were seeded in 96-well plates, and once each time point finished, culture medium was removed and the cells incubated at room temperature for 20 min in 0.5% crystal violet 20% methanol staining solution. Next, the plates were washed carefully four times in a gentle stream of tap water, inverted on a filter paper to remove the remaining liquid, and dried at room temperature for at least 1 h. Finally, crystal violet was solubilized by adding 200 μL of methanol per well. OD_570_ was measured using the Infinite 200 PRO plate reader (Tecan™).

### Measurement of cytoplasmic and mitochondrial Ca^2+^ signals

Cells were grown on 18 mm Ø glass coverslips in 6-well plates and cytoplasmic and mitochondrial Ca^2+^ signals were evaluated by *timelapse* in confocal microscopy, using Nikon C^2+^ Confocal Microscope System (Nikon™), or Leica TCS SP8 Confocal Laser Scanning Microscope (Leica™) with control of temperature (37° C), humidity and CO_2_ (5%).

Measurements were performed in calcium-free medium, using DMEM no calcium (Gibco™ #21068) supplemented with 100 µM EGTA.

*Cytoplasmic Ca^2+^ signals*: IP_3_R activity was evaluated using the Ca^2+^ sensitive cytoplasmic probe Fluo-4/AM (Max. ex/em: 494/506 nm). IMR90 cells growth in coverslips were loaded with 5 µM Fluo-4/AM (30 min) in complete medium and mounted in round chamber for microscopy imaging. The cells were washed twice with calcium-free medium and imaged in presence of the corresponding compounds. After a period of baseline recording, the cells were stimulated with a pulse of 0,3 mM histamine to induce IP_3_R-mediated Ca^2+^ release. A total of 140 frames were recorded every 1.5 seconds approximately, at 488 nm excitation and using a 63x objective. Images were analyzed and quantified using ImageJ (NIH).

*Mitochondrial* Ca^2+^ *signals*: Mitochondrial Ca^2+^ uptake from IP_3_R-mediated Ca^2+^ release after histamine stimuli was determined initially using the mitochondria-targeted genetically encoded fluorescent Ca^2+^ indicator CEPIA2mt (K_d_=160 nM; Max. ex/em: 488/500 and 550 nm). For these experiments, cells were transfected with plasmid DNA (2 µg/well on 6-well plates) using lipofectamine 3000. After 24 h pos-transfection, cells were trypsinized, seeded on 25 mm Ø glass coverslips and finally cultured for 24 h. Confocal image recording was performed as described above, at 488 nm excitation laser. In addition, we used Rhod-2/AM (Max. ex/em: 552/581 nm) to measure mitochondrial Ca^2+^ signals, co-localized with the mitotracker green FM (Max. ex/em: 490/516 nm) labeling. For this approach, cells seeded on 25 mm Ø glass coverslips were loaded with 5 µM Rhod-2/AM and 100 nM mitotracker green FM (30 min) in complete medium after completion of the experimental treatments. For the confocal timelapse, cells were mounted in the microscope chambers, washed twice with calcium-free medium, and finally recorded before and after the stimulation with 0,3 mM histamine, using a 63x objective, at 488 nm and 561 nm excitation lasers for mitotracker green FM and Rhod-2/AM respectively. In addition, we used an FCCP stimuli to uncouple mitochondrial membrane potential, after a period of baseline recording. This resulted in the release of mitochondrial Ca^2+^, which was measured in the cytoplasm using the Fluo-4/AM probe. IMR90 cells were loaded with 5 µM Fluo-4/AM for 30 minutes in complete medium, as previously described. A total of 140 frames were recorded every 1.5 seconds approximately. Images were analyzed and quantified using ImageJ (NIH).

*Spontaneous Ca^2+^ activity:* IMR90 Fibroblasts were loaded with Fluo-4 AM (2 µM) and incubated for 30 min in complete darkness at 37 °C. After incubation, the cells were washed twice with 1× PBS to remove excess dye and subsequently maintained in FluoroBrite™ DMEM for live-cell imaging. Ca^2+^ dynamics were recorded using a Zeiss Airyscan confocal microscope equipped with a Plan-Apochromat 40× objective, 1 Airy unit pinhole, and 16-bit acquisition (1772 × 1772 pixels). The pixel dwell time was set to 1 µs, with laser power at 3%. Time-lapse imaging was performed as a series of 100–150 frames acquired at 10 frames per second, under temperature-controlled conditions (37 °C).

*ER Ca^2+^ measurement*: IMR90 were transfected with G-CEPIAer. Forty-eight hours post transfection cells were recorded. The protocol consisted of basal recording with Ringer’s modified with 2 mM CaCl2, 7 min of reticular Ca^2+^ depletion using Ca^2+^-free Ringer’s with 300 µM histamine to finally replace the milieu with modified Ringer’s 2 mM CaCl2 solution. Images were acquired every 5 s for 200 frames using a TIRFM 60× objective (N.A. = 1.45) in an Olympus IX71 microscope equipped with an FLIR camera (Backfly S, BFS-U3-51S5M, FLIR, Richmond, BC, Canada). Fluorescence intensity was quantified using the Image J2/FIJI v.2.9.0 software (National Institute of Mental Health, Bethesda, MD, USA) and data were normalized using the initial ratio of fluorescence.

### Protein extraction and quantification

Cells seeded in 6-well plates were cultured for 24 h and then subjected to corresponding experimental treatments. Once treatments were finished, cells were placed on ice, washed twice with ice-cold PBS, and finally scraped and lysed using CytoBuster™ protein extraction reagent supplemented with protease and phosphatase inhibitors. The obtained homogenates were sonicated, incubated 15 minutes on ice, centrifugated at 12,500 x *rpm* for 20 minutes at 4 °C, and the supernatant collected. Protein quantification was performed using the Bradford method in 96-well plates and OD_595_ nm was determined using the Infinite 200 PRO plate reader (Tecan™).

### SDS-PAGE and Western Blotting

Protein extracts were mixed with sample buffer 5x (250 mM Tris-HCl; 40% v/v glycerol; 8% v/v 2-mercaptoethanol; 10% w/v SDS; 0,5% w/v bromophenol blue, pH 6.8) and then boiled for 5 minutes at 100 °C. Once cooled, 30 µg of protein extract from each sample were separated electrophoretically in either 5%, 10% or 15% SDS-polyacrylamide gels using a Mini-PROTEAN Tetra Cell (Bio-Rad™) with running buffer containing: 25 mM Tris; 192 mM glycine; 0,1% w/v SDS at pH 8.3, and subsequently transferred to PDVF membranes by Trans-Blot™ SD Semi-Dry Transfer Cell (Bio-Rad ®). Membranes were subsequently blocked 60 minutes with 5% w/v bovine serum albumin (BSA) prepared in Tris buffered saline-tween (TBS-T) containing: 20 mM Tris-HCl, 150 mM NaCl and 0.05% w/v Tween 20, at pH 7.5. Corresponding primary antibodies (dilution 1:1000 or 1:3000) were incubated overnight at 4 °C, then washed with TBS-T and subsequently incubated with HRP-conjugated secondary antibody (dilution 1:2000) for 1 h at room temperature with shaking. Protein signal was visualized using SuperSignal™ West Pico PLUS chemiluminescent substrate and documented with ChemiDoc™ Imaging System (Bio-Rad). Images were analyzed and quantified using ImageJ (NIH).

### Plasma Membrane Proteomic Screening

Plasma membrane proteomic screening was performed essentially as described^53^.

### Immunofluorescence and PLA

For Immunofluorescence and PLA assays, 5,000 cells per cm^2^ were seeded on 12 mm Ø glass coverslips. Once the corresponding experimental treatments were completed, cells were fixed with 4% w/v paraformaldehyde (Image-iT™ fixative solution; Thermo Fisher Scientific), washed in PBS, then permeabilized in 0.1% v/v triton X-100 in PBS and blocked for 1 h in 10% w/v BSA in PBS at room temperature. After blocking, cells were incubated with the indicated antibodies (also for PLA): anti-Bcl-xL(54H6), anti-VDAC1 (AB10527), anti- VDAC1 (AB14734), anti-TOMM20 (AB56783); overnight at 4 °C followed by either staining with Alexa-conjugated secondary antibodies (Molecular Probes) for 2 h at room temperature and protected from light or following Duolink manufacturer’s instructions (Duolink, Sigma-Aldrich). Once incubation was completed, cells were washed 3 times in PBS and Hoechst 33342 was applied during the last wash. The coverslips were mounted on slides using the mounting medium Fluoromount-G™. Images were obtained using a confocal microscope (TCS SP8 Spectral Confocal Microscope; Leica) and subsequently processed and analyzed using ImageJ software (NIH). For the analysis of γ-H2A.X, the number of foci (positive dots) within each nucleus was quantified

### Molecular docking and Structural Comparison

Molecular docking was initially performed using ColabFold^54^ , and the best structure was selected based on established scoring metrics and structural parameters. The crystal structures used as templates were PDB ID: 3EMN for VDAC1 and PDB ID: 4QNQ for Bcl- xL. All predicted conformations converged toward a common binding mode, indicating a consistent interaction interface. The selected conformation was then used as input to generate 5,000 interaction models with the Rosetta software package^55^. The best conformer was chosen according to the IC_Score and subsequently analyzed using PyMOL (Schrödinger, LLC). In addition, a structural comparison was performed between the docking-derived VDAC1–Bcl-xL complex and the crystal structure of Bcl-xL bound to ABT- 263 (PDB ID: 4QNQ), to evaluate potential overlap at the binding site.

### Mice and doxorubicin-induced *in vivo* senescence

This study was carried following the strict recommendations in the Guide for the Care and Use of Laboratory Animals of the National Institutes of Health. The animal protocol was approved by the Committee on the Ethics of Animal Experiments of Universidad Mayor (number:05/2020). Mice were provided standard chow ad libitum and maintained under a 12:12-hour light/dark cycle. Systemic cellular senescence was introduced by treating mice with doxorubicin, as described^56^. Briefly, a single dose of doxorubicin (10 mg/kg) was injected intraperitoneally and/or ABT-263 inhibitor was administered by gavage. Organs (lung, kidney, and liver) were collected. To avoid variations due to circadian cycle, all animals were sacrificed between 10 am and 2 pm^57^.

### Statistical analysis

Depending on the type of experiment, the results are presented as representative images or as the mean ± SEM of at least three independent experiments. Statistical analyses were performed using Prism 8 software (GraphPad software). Significance of differences was assessed using unpaired t-tests. In the case of data that exhibited a normal distribution, the data were analyzed using a T-Test and one-way ANOVA with comparisons among the different experimental groups using Sidak correction. Additionally, comparisons were made with the control group using Dunnett’s test for multiple comparisons. Differences with a *p* value <0.05 were considered statistically significant.

---

**Supplementary Figure S1.**
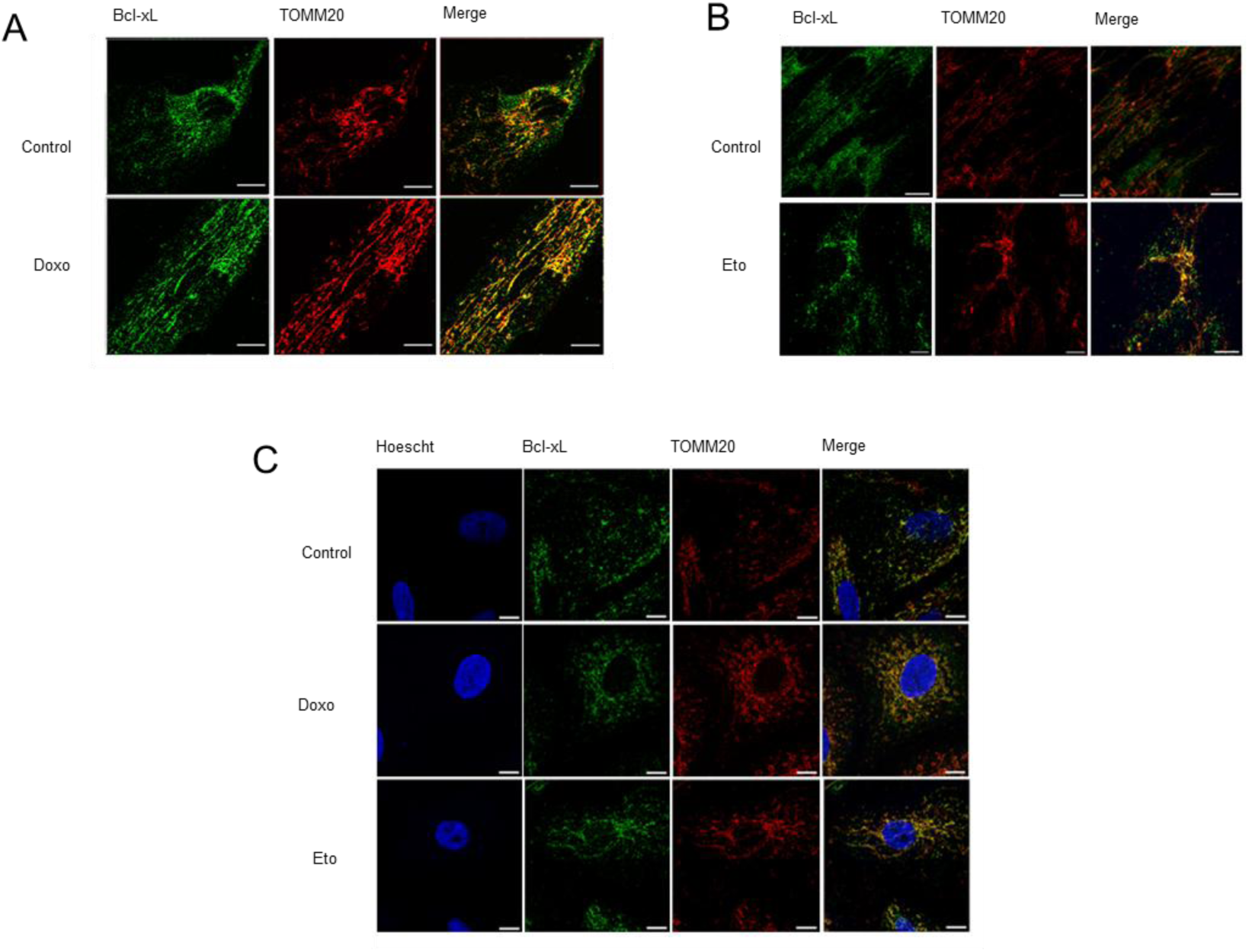
Validation of Bcl-xL mitochondrial localization in early senescent cells. (A–B) Representative immunofluorescence images showing colocalization of Bcl-xL (green) with the mitochondrial marker TOMM20 (red), and their corresponding merged images, in IMR90 fibroblasts treated with either 250 nM Doxorubicin (A) or 40 µM Etoposide (B) for 24 hours. (C) Representative immunofluorescence images of A549 cells treated with either 250 nM Doxorubicin or 40 µM Etoposide for 24 hours, showing colocalization of Bcl-xL (green) with TOMM20 (red). Nuclei were stained with Hoechst (blue). Merged images indicate overlapping signals consistent with mitochondrial localization of Bcl-xL. Images shown are representative of three independent experiments per condition. Scale bar: 10 μm.

**Supplementary Figure S2.**
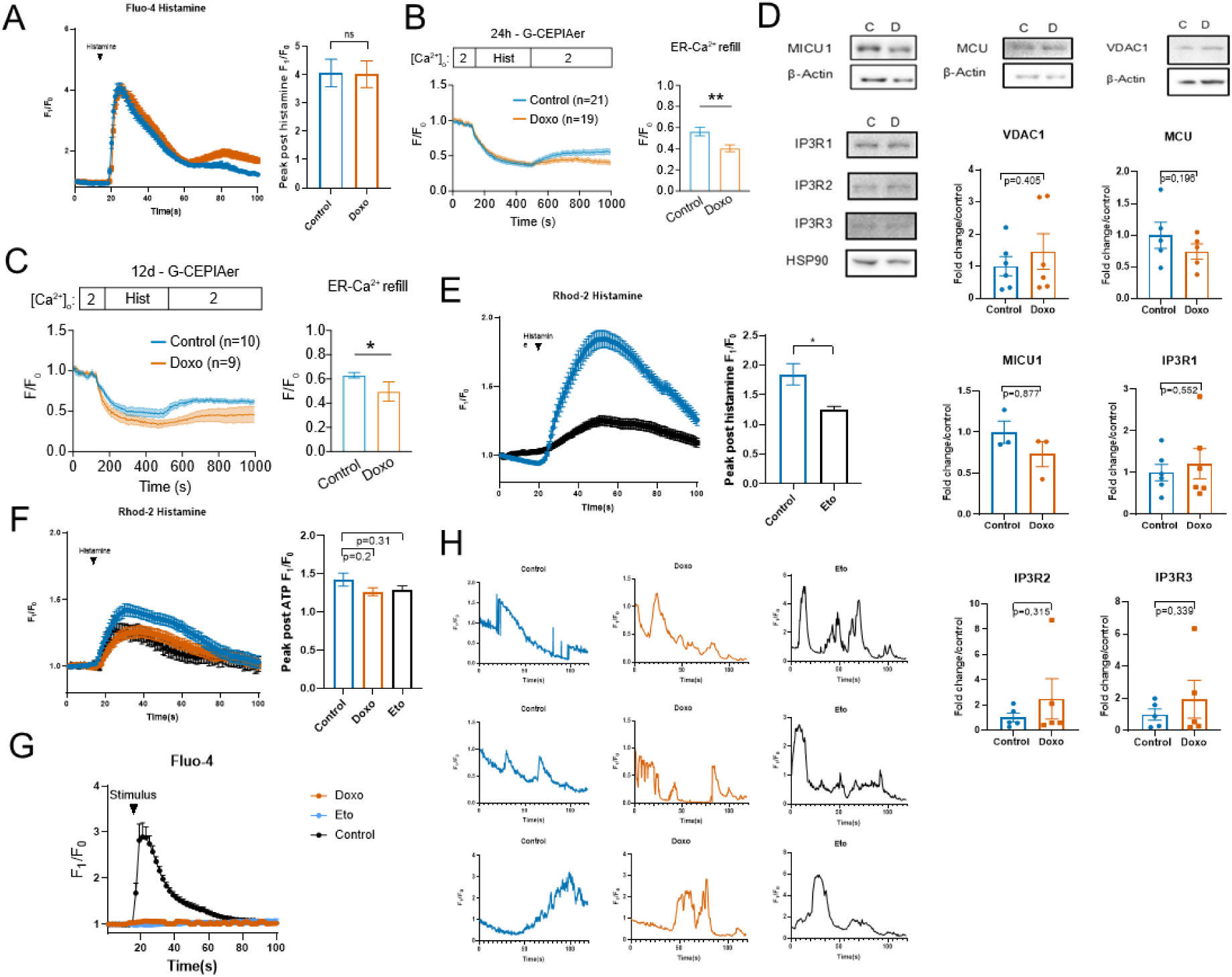
**Mitochondrial and ER Ca²**⁺ **dynamics during therapy-induced senescence (TIS).** (A) Cytosolic Ca²⁺ transients measured using Fluo-4 AM in IMR90 fibroblasts treated with 250 nM Doxorubicin, showing no significant differences in peak response to ATP stimulation compared to control(B) Representative normalized fluorescence traces (F/F0) of the ER Ca^2+^ indicator G-CEPIA1er showing luminal Ca^2+^ dynamics following histamine stimulation (300 µM) under 2 mM [Ca^2+^]0. Traces correspond to growing IMR90 cells (blue, n=10) and IMR90 cells treated with doxorubicin for 24h (red, n=9). The transient drop reflects histamine-induced ER Ca²⁺ release followed by the recovery (refill) phase. (C) Quantification of ER Ca^2+^ refill expressed as F/F₀ at the end of the recording. Data are presented as mean ± SD. A significant reduction in ER Ca²⁺ refill was observed in doxorubicin-treated cells compared to growing controls (**p < 0.01, two-tailed Mann–Whitney test). (D) Western blot analysis and quantification of calcium-handling proteins in IMR90 fibroblasts under control (C) or Doxorubicin-treated (D) conditions. Assessed components include VDAC1, MCU, MICU1, and the IP3 receptor isoforms (IP3R1, IP3R2, IP3R3). Protein levels were normalized to β-actin or HSP90 (for IP3Rs). (E) Mitochondrial Ca²⁺ uptake in IMR90 cells assessed with Rhod-2 AM following stimulation. Etoposide-treated cells exhibited reduced mitochondrial Ca²⁺ uptake compared to control, as shown by the F₀/F₁ peak ratio: Control: 1.845 ± 0.31; Eto: 1.25 ± 0.08 (F) Mitochondrial Ca²⁺ uptake in A549 cells assessed with Rhod-2 AM following stimulation. Both Doxorubicin- and Etoposide-treated cells exhibited reduced mitochondrial Ca²⁺ uptake compared to control, as shown by the F₀/F₁ peak ratio: Control: 1.423 ± 0.08487; Doxo: 1.264 ± 0.05223; Eto: 1.290 ± 0.05196. (G) Cytosolic Ca²⁺ responses to doxo, eto and histamine stimulation measured by Fluo-4, showing no significant differences across treatments. (H) Representative Ca^2+^ traces showing fluorescence ratio signals (F_1_/F0) of IMR90 cells recorder one hour after being stimulated with either 250nM doxorubicin (Doxo), 40 µM etoposide (Eto) or vehicle (control) in normal media (N=3). Data are presented as mean ± SEM from at least three independent experiments per condition.

**Supplementary Figure S3.**
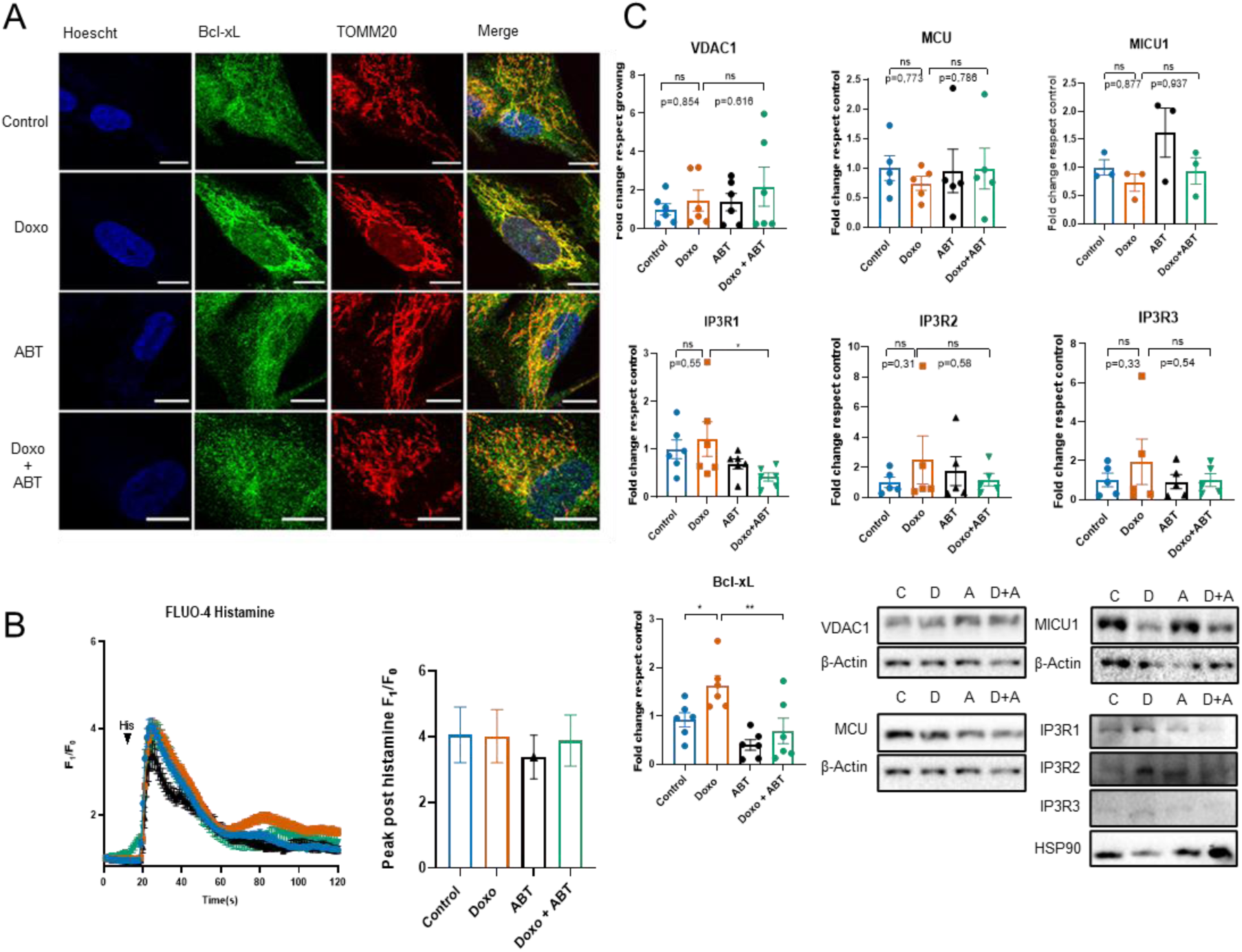
**ABT-263 disrupts Bcl-xL localization and affects mitochondrial Ca²**⁺ **uptake in senescent cells.** (A) Immunofluorescence images showing decreased colocalization between Bcl-xL (green) and TOMM20 (red) in IMR90 cells treated with Doxorubicin (250 nM), ABT-263 (3 µM), or their combination for 24 hours. Nuclei stained with Hoechst (blue). (B) Western blot quantification of VDAC1, MCU, MICU1, IP3Rs, and Bcl-xL protein levels under the same conditions, normalized to β-actin or HSP90. (C) Cytosolic Ca²⁺ responses to histamine stimulation measured by Fluo-4, showing no significant differences across treatments. Quantitative analyses represent mean ± SEM. Scale bar: 10 μm. (at least three independent experiments per condition). All panels: one- way ANOVA.

**Supplementary Figure S4.**
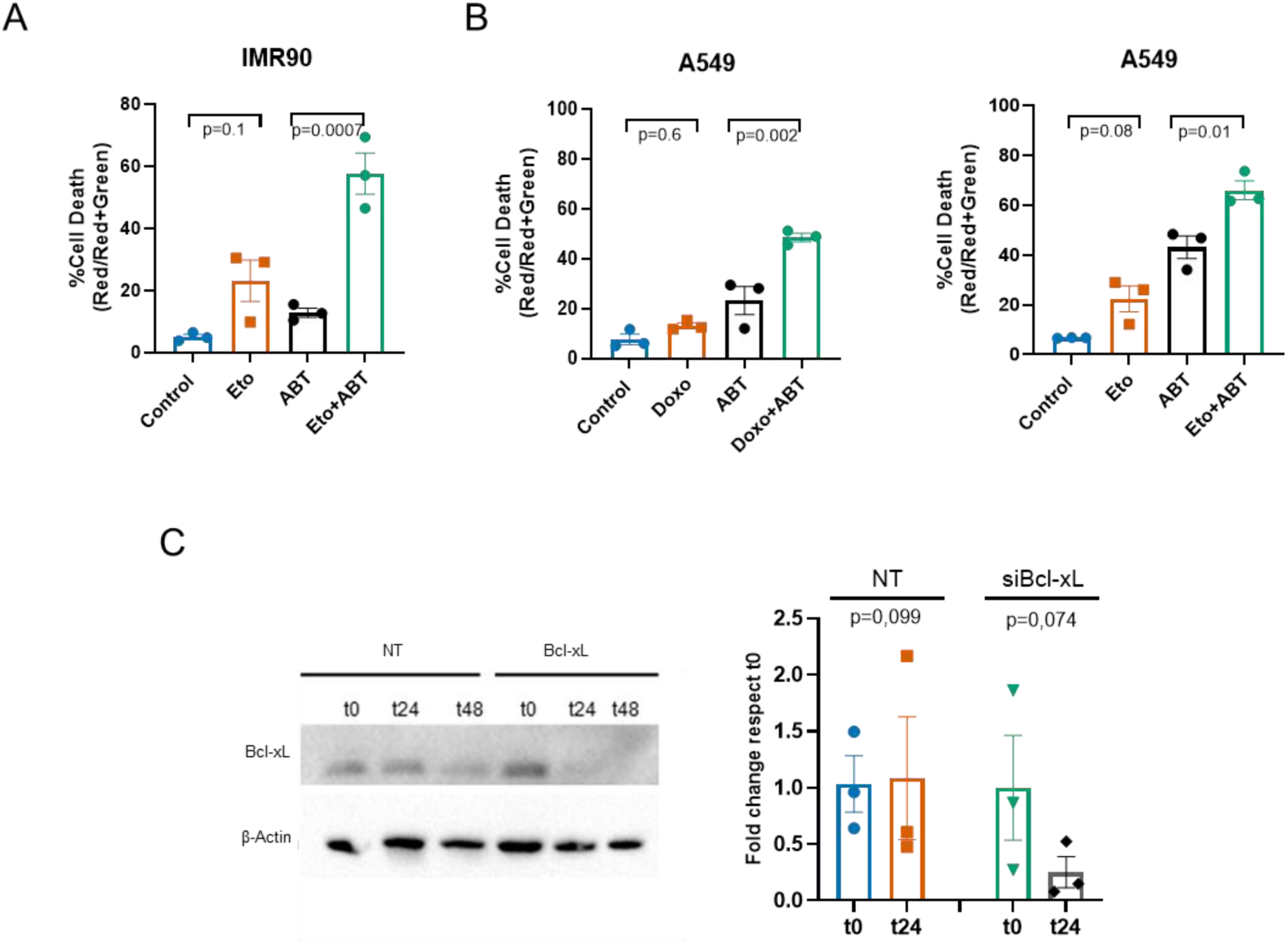
Genetic validation of Bcl-xL knockdown and ABT- 263-induced apoptosis. (A) Quantification of cell death using Sytox Green in IMR90 fibroblasts treated with Etoposide (10 µM), ABT-263 (3 µM), or the combination. ABT-263 significantly enhanced cell death when combined with Etoposide compared to either treatment alone (Control: 5.033 ± 0.7881; Eto: 23.13 ± 6.679; ABT: 12.83 ± 1.477; Eto + ABT: 57.67 ± 6.617). (B) Sytox Green quantification of cell death in A549 cells treated with either 40 µM Etoposide or 250 nM Doxorubicin, with or without ABT-263 (3 µM). In both cases, ABT-263 significantly increased cell death when used in combination. Doxorubicin-treated cells: Control: 7.700 ± 2.066; Doxo: 13.23 ± 1.102; ABT: 23.27 ± 5.596; Doxo + ABT: 48.63 ± 1.656. Etoposide-treated cells: Control: 6.767 ± 0.08819; Eto: 22.43 ± 5.185; ABT: 43.13 ± 4.537; Eto + ABT: 66.03 ± 3.840. (C) Western blot analysis confirming efficient knockdown of Bcl-xL in IMR90 fibroblasts at 24- and 48-hours post- transfection with Bcl-xL siRNA. Protein levels were normalized to β-actin and expressed as fold change relative to respective time 0. Non-targeting control: t0: 1.033 ± 0.2496; t24: 1.084 ± 0.5433. Bcl-xL siRNA: t0: 1.000 ± 0.4636; t24: 0.2513 ± 0.1376. Data are presented as mean ± SEM from at least three independent experiments per condition. Panels S4A– S4B: one-way ANOVA. S4C: T-Test.

